# Identification and Functional Characterization of Up-Regulated Hub Genes in Adenocarcinoma Across Multiple Organ Sites

**DOI:** 10.1101/2024.08.21.609070

**Authors:** Ricardo Romero, Dulce M. A. Bastida

## Abstract

Adenocarcinoma is a prevalent and aggressive form of cancer affecting various organ systems. To gain deeper insights into the molecular mechanisms underlying adenocarcinoma, we conducted a comprehensive bioinformatics analysis to identify consistently up-regulated hub genes and their regulatory networks across multiple adenocarcinoma types. By integrating gene expression data from GEO and GEPIA2 databases, we identified 129 commonly up-regulated genes, from which 10 key hub genes were further characterized: CHEK1, CDC20, ANLN, RRM2, CCNB1, CCNA2, KIF23, TOP2A, BUB1, and KIF11. These hub genes were found to be significantly overexpressed in several adenocarcinoma types, including colorectal, lung, pancreatic, and ovarian cancer, except for prostate adenocarcinoma. The regulatory network analysis revealed that the expression of these hub genes is controlled by transcription factors, such as YBX1, E2F1, MYC, E2F3, and TP53, as well as miRNAs, including hsa-miR-103a-3p, hsa-let-7e-5p, and hsa-miR-15b-5p, which were consistently downregulated across the adenocarcinoma types studied. Functional enrichment analysis highlighted the involvement of these hub genes in critical cellular processes, including cell cycle regulation, mitotic division, and cellular senescence. These findings provide a comprehensive understanding of the molecular landscape of adenocarcinoma and identify potential therapeutic targets and prognostic biomarkers for further investigation.

## 1 Introduction

Adenocarcinoma, a malignant tumor originating in glandular epithelial cells, is a prevalent form of cancer affecting various organs, including the lungs, colon, pancreas, and prostate [1]. This type of cancer accounts for approximately 40% of all lung cancers and is the most common histological subtype of colorectal cancer [2, 3]. The incidence of adenocarcinoma has been steadily increasing worldwide, posing a significant global health challenge [4].

The molecular pathogenesis of adenocarcinoma involves complex genetic and epigenetic alterations that disrupt normal cellular processes, leading to uncontrolled cell growth and division [5]. Key molecular events in adenocarcinoma development include mutations in oncogenes such as KRAS, EGFR, and BRAF, as well as inactivation of tumor suppressor genes like TP53 and CDKN2A [6, 7]. These genetic alterations result in the dysregulation of critical signaling pathways, including the MAPK/ERK, PI3K/AKT, and Wnt/β-catenin pathways, which contribute to tumor initiation, progression, and metastasis [8].

Advances in molecular profiling techniques, particularly next-generation sequencing, have significantly enhanced our understanding of the genomic landscape of adenocarcinoma [9]. This knowledge has led to the identification of numerous potential therapeutic targets and the development of targeted therapies. For instance, EGFR tyrosine kinase inhibitors (TKIs) have shown remarkable efficacy in treating EGFR-mutant lung adenocarcinoma, while BRAF inhibitors have demonstrated promising results in BRAF V600E-mutant colorectal adenocarcinoma [10, 11].

Despite these advancements, several challenges remain in the management of adenocarcinoma. First, the heterogeneity of these tumors, both inter- and intra-tumorally, complicates treatment strategies and contributes to drug resistance [12]. Second, while targeted therapies have shown initial success, many patients eventually develop resistance, necessitating the identification of new therapeutic targets and combination strategies [13]. Third, early detection of adenocarcinoma remains challenging, particularly for organs such as the pancreas, where symptoms often appear only in advanced stages [14].

Furthermore, our understanding of the complex interplay between genetic alterations, epigenetic modifications, and the tumor microenvironment in adenocarcinoma progression is still incomplete [15]. Additionally, the role of non-coding RNAs, such as long non-coding RNAs (lncRNAs) and microRNAs (miRNAs), in regulating gene expression and contributing to adenocarcinoma development is an emerging area of research that requires further investigation [16].

To address these challenges and improve patient outcomes, there is a critical need for comprehensive gene expression profiling of adenocarcinoma across different organ systems. Such studies can potentially reveal novel biomarkers for early detection, identify new therapeutic targets, and provide insights into the mechanisms of drug resistance. Moreover, integrating gene expression data with other omics data, such as proteomics and metabolomics, could offer a more holistic understanding of adenocarcinoma biology and guide personalized treatment approaches [17].

In this study, we aim to analyze the gene expression profiles of adenocarcinoma, from multiple organ sites, using a comprehensive bioinformatics approach. There is a growing recognition of the need for comprehensive genomic and transcriptomic profiling to uncover the full spectrum of genetic alterations and gene expression changes in adenocarcinoma. Such efforts are essential to address the gaps in our understanding of the disease and to develop more effective, personalized treatment strategies. This paper aims to contribute to this effort by investigating the landscape of expressed genes in adenocarcinoma, with the goal of identifying new targets for therapy and improving the clinical management of the disease.

## 2 Methods

Our study employed a comprehensive bioinformatics approach to identify and analyze up-regulated hub genes in adenocarcinoma across multiple organ sites. The methodology consisted of several key steps:

### Data Collection and Differential Gene Expression Analysis

To identify up-regulated hub genes in adenocarcinoma, we conducted a differential gene expression analysis using datasets from the Gene Expression Omnibus (GEO) and GEPIA2 databases. Three adenocarcinoma studies from GEO—GSE32863,[18] GSE10972,[19] and GSE15471 [20, 21]—were selected for analysis. The criteria for identifying up-regulated genes were set to a log fold change (logFC) greater than 0 and an adjusted p-value less than 0.05. The FunRich software (http://www.funrich.org/) was used to identify genes commonly up-regulated across all three datasets.

In parallel, we performed a differential gene expression analysis using GEPIA2 [22] across five different types of adenocarcinoma: colon (COAD), lung (LUAD), pancreas (PAAD), rectum (READ), and stomach (STAD). For this analysis, the threshold was set to a logFC greater than 1 and an adjusted p-value less than 0.05. The Venn diagram web tool from https://bioinformatics.psb.ugent.be/webtools/Venn/ was employed to identify genes commonly up-regulated across all five adenocarcinoma types.

### Integration and Protein-Protein Interaction (PPI) Network Construction

The intersection of the gene sets obtained from the GEO and GEPIA2 analyses was determined using FunRich. The intersecting genes were then used to construct a protein-protein interaction (PPI) network via StringDB [23], setting a high confidence threshold of 0.7 to ensure robust interactions. The resulting PPI network was imported into Cytoscape [24] for further analysis.

### Hub Gene Identification

Within Cytoscape, the CytoHubba [25] plug-in was utilized to identify the top 10 hub genes based on two key metrics: degree and betweenness centrality. This dual approach of using both degree and betweenness centrality ensured that the hub genes identified were not only highly connected but also central to the overall network structure, reflecting their potential significance in adenocarcinoma.

To validate the expression levels of the identified hub genes in adenocarcinoma, we conducted a secondary verification using the GEPIA2 database. This step confirmed the consistency of the hub genes’ up-regulation across multiple datasets and provided further support for their relevance in adenocarcinoma.

### miRNA and Transcription Factor Target Prediction

The validated hub genes were then submitted to MirWalk [26] to predict miRNAs targeting these genes. To enhance the reliability of the predictions, only miRNAs that were validated by three major databases—TargetScan [27], miRDB [28], and miRTarBase [29]—were considered. The expression levels of the identified miRNAs across different cancer types were further analyzed using the OncoMir database. [30]

In addition, transcription factors (TFs) targeting the hub genes were identified using the TRRUST database.[31] This step provided insights into the regulatory mechanisms governing the expression of the hub genes in adenocarcinoma.

### Functional enrichment analysis

Hub genes and TFs were submitted to GeneTrail [32] to perform functional enrichment analysis, focusing on molecular function and biological processes from the Gene Ontology knowledgebase,[33, 34] as well as KEGG (Kyoto Encyclopedia of Genes and Genomes) pathways.

### Construction of the Regulatory Network

Finally, we integrated the hub genes, their targeting miRNAs, and transcription factors into a comprehensive regulatory network. This network was constructed using StringDB and visualized in Cytoscape, providing a detailed map of the molecular interactions and regulatory pathways involved in adenocarcinoma.

This methodological approach ensured the identification of the most consistently up-regulated hub genes in adenocarcinoma, along with their associated regulatory elements, providing a robust framework for further investigation into their roles in cancer development and progression.

## 3 Results

### Identification of Up-Regulated Genes in Adenocarcinoma

Through differential gene expression analysis, we identified 875 up-regulated genes that were common across the three GEO datasets (GSE32863, GSE10972, and GSE15471, Fig. 1). Similarly, in the GEPIA2 analysis, 471 up-regulated genes were found to be common across five adenocarcinoma types (COAD, LUAD, PAAD, READ, and STAD, Fig. 2). The intersection of these two gene sets yielded 129 genes that were consistently up-regulated across both sources (Fig. 3).

**Fig. 1:**
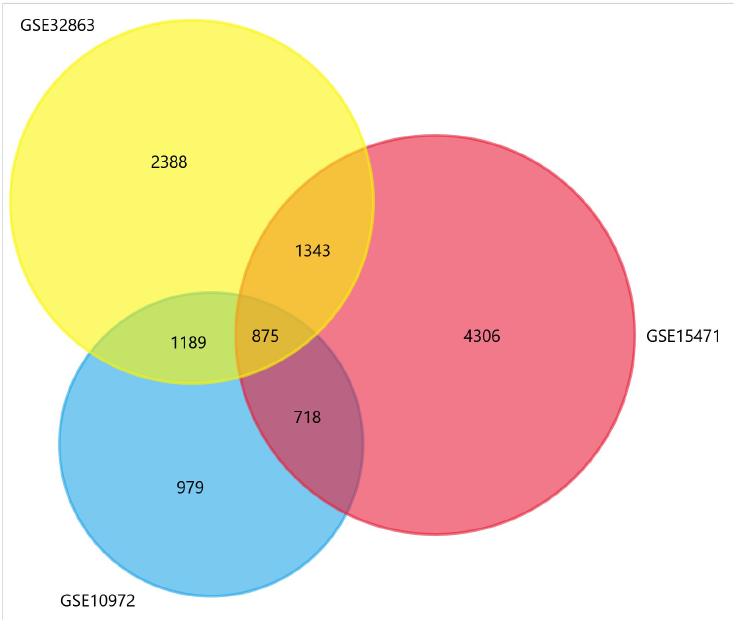
Venn Diagram of Up-Regulated Genes Identified Across Three Adenocarcinoma GEO Datasets. This Venn diagram shows the overlap of up-regulated genes identified in three different GEO datasets: GSE32863 (yellow), GSE10972 (blue), and GSE15471 (red). The numbers within each section represent the number of up-regulated genes unique to or shared among the datasets. - **GSE32863**: 2,388 up-regulated genes are unique to this dataset. -**GSE10972**: 979 up-regulated genes are unique to this dataset. -**GSE15471**: 4,306 up-regulated genes are unique to this dataset. -**Common Across All Three Datasets**: 875 genes are commonly up-regulated in all three datasets. This diagram illustrates the intersection of gene expression profiles across different adenocarcinoma studies, highlighting the subset of 875 genes that are consistently up-regulated across all datasets.

**Fig. 2:**
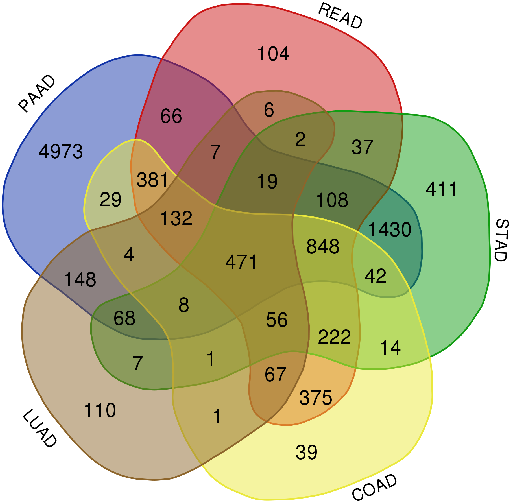
Venn Diagram of Up-Regulated Genes Across Five Adenocarcinoma Types Using GEPIA2 Data. This Venn diagram displays the overlap of up-regulated genes identified across five different types of adenocarcinomas analyzed using the GEPIA2 platform: PAAD (pancreatic adenocarcinoma), READ (rectum adenocarcinoma), STAD (stomach adenocarcinoma), COAD (colon adenocarcinoma), and LUAD (lung adenocarcinoma). -**PAAD**: 4,973 up-regulated genes are unique to pancreatic adenocarcinoma. **-READ**: 104 up-regulated genes are unique to rectum adenocarcinoma. -**STAD**: 411 up-regulated genes are unique to stomach adenocarcinoma. -**COAD**: 375 up-regulated genes are unique to colon adenocarcinoma. -**LUAD**: 110 up-regulated genes are unique to lung adenocarcinoma. -**Common Across All Five Types**: 471 genes are commonly up-regulated in all five types of adenocarcinomas, suggesting their potential role as core drivers of adenocarcinoma pathology.The diagram illustrates the complex overlap between the gene expression profiles of these adenocarcinoma types, highlighting both unique and shared molecular characteristics.

**Fig. 3:**
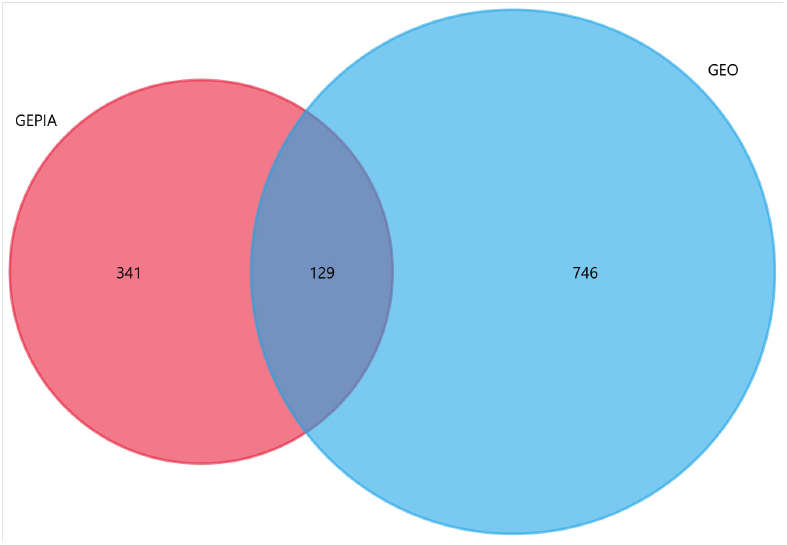
Venn Diagram Showing the Overlap of Up-Regulated Genes Identified in GEPIA2 and GEO Datasets. This Venn diagram illustrates the overlap of up-regulated genes identified from two different data sources: GEPIA2 (red) and GEO (blue). -**GEPIA2**: A total of 341 up-regulated genes were uniquely identified using the GEPIA2 platform. -**GEO**: A total of 746 up-regulated genes were uniquely identified across the selected GEO datasets. -**Common Genes**: The intersection of the two sets reveals 129 genes that are commonly up-regulated in both GEPIA2 and GEO datasets. This diagram highlights the consistency and differences between the two data sources, identifying a core set of 129 up-regulated genes that are robustly associated with adenocarcinoma across both platforms. These common genes are likely to represent key molecular drivers of adenocarcinoma and are prioritized for further analysis in the study.

### Construction and Analysis of the PPI Network

The protein-protein interaction (PPI) network constructed from these 129 common genes included 500 edges and 129 nodes, with an average degree of 7.75 and an average local clustering coefficient of 0.41. The network exhibited a significant PPI enrichment P-value of less than 1.0e-16, indicating that the identified genes are more interconnected than would be expected by chance, suggesting a high level of functional association among them.

### Identification of Hub Genes

Using CytoHubba in Cytoscape, we identified 10 hub genes within the PPI network based on degree and betweenness centrality metrics: CHEK1, CDC20, ANLN, RRM2, CCNB1, CCNA2, KIF23, TOP2A, BUB1, and KIF11. These hub genes were found to be significantly overexpressed in six adenocarcinoma types (COAD, LUAD, PAAD, READ, STAD, and ovarian serous cystadenocarcinoma, OV). However, in prostate adenocarcinoma (PRAD), the genes KIF11, KIF23, CCNA2, ANLN, and BUB1 did not exhibit significant overexpression. (Fig. 4, we present here the general heat map, detailed boxplots for each gene, with significance annotations, are provided as supplementary material).

**Fig. 4:**
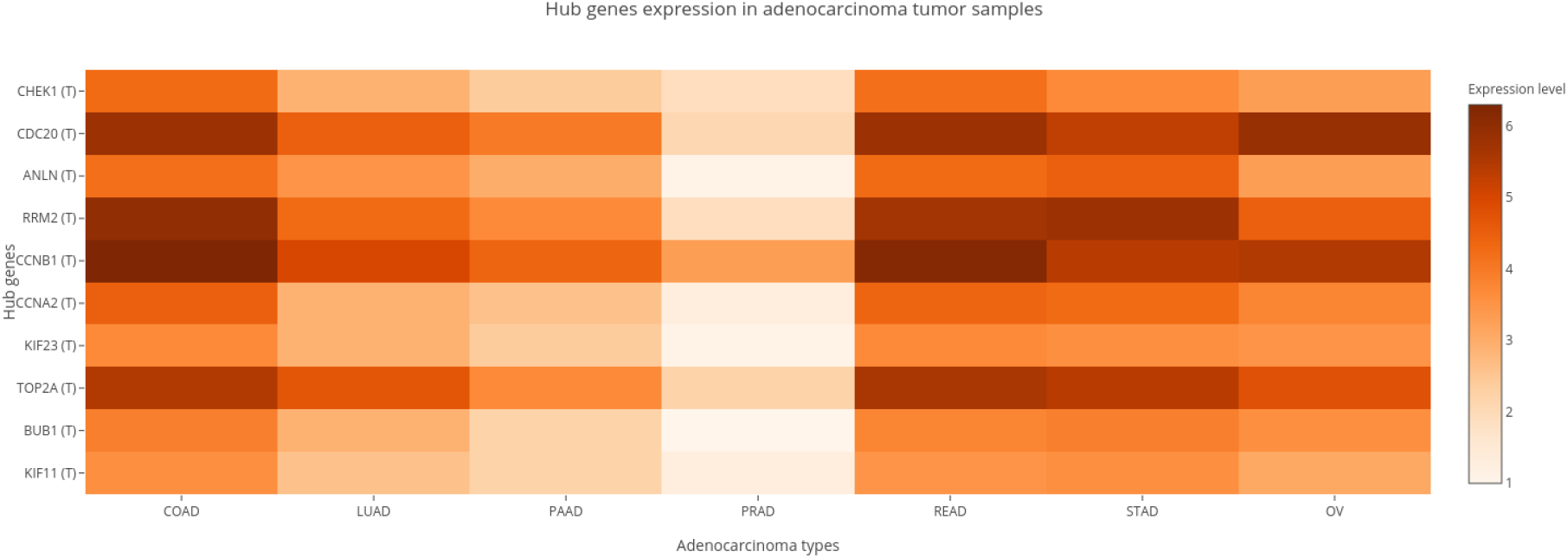
Heatmap of Hub Gene Expression Levels Across Different Adenocarcinoma Types. This heatmap illustrates the expression levels of key hub genes identified in adenocarcinoma across various tumor samples. The hub genes are listed on the y-axis, including CHEK1, CDC20, ANLN, RRM2, CCNB1, CCNA2, KIF23, TOP2A, BUB1, and KIF11. The different types of adenocarcinomas are represented on the x-axis: COAD (colon adenocarcinoma), LUAD (lung adenocarcinoma), PAAD (pancreatic adenocarcinoma), PRAD (prostate adenocarcinoma), READ (rectum adenocarcinoma), STAD (stomach adenocarcinoma), and OV (ovarian serous cystadenocarcinoma). - **Color Gradient**: The color gradient represents the expression levels of these genes, with darker shades of orange and brown indicating higher expression levels, and lighter shades indicating lower expression levels. - **Key Insights**: The heatmap reveals that most of these hub genes are consistently overexpressed across the various adenocarcinoma types, with some variability in expression levels. Notably, PRAD (prostate adenocarcinoma) exhibits lower expression levels for certain hub genes such as CCNA2, KIF23, and BUB1, which aligns with previous findings indicating that these genes are not significantly overexpressed in PRAD compared to other adenocarcinoma types.This visualization highlights the potential significance of these hub genes in the progression of adenocarcinoma and underscores their relevance as potential therapeutic targets or biomarkers across different adenocarcinoma subtypes.

### Regulatory Network Involving Transcription Factors and miRNAs

We further explored the regulatory landscape of these hub genes by identifying key transcription factors and miRNAs that target them. The transcription factors identified include YBX1, E2F1, MYC, E2F3, and TP53. The miRNAs targeting these hub genes include hsa-miR-503-5p, hsa-miR-106b-5p, hsa-miR-16-5p, hsa-miR-103a-3p, hsa-let-7b-5p, hsa-let-7c-5p, hsa-let-7d-5p, hsa-let-7e-5p, hsa-let-7f-5p, hsa-let-7i-5p, hsa-miR-15b-5p, hsa-miR-195-5p, hsa-miR-424-5p, and hsa-miR-497-5p. Among these, miRNAs hsa-miR-103a-3p, hsa-let-7e-5p, and hsa-miR-15b-5p were consistently down-regulated across all adenocarcinoma types in the Oncomir database (COAD, READ, LUAD, STAD, PRAD, and PAAD) (Fig. 5). The integrated regulatory network comprising hub genes, TFs, and miRNAs is presented in Fig. 6.

**Fig. 5:**
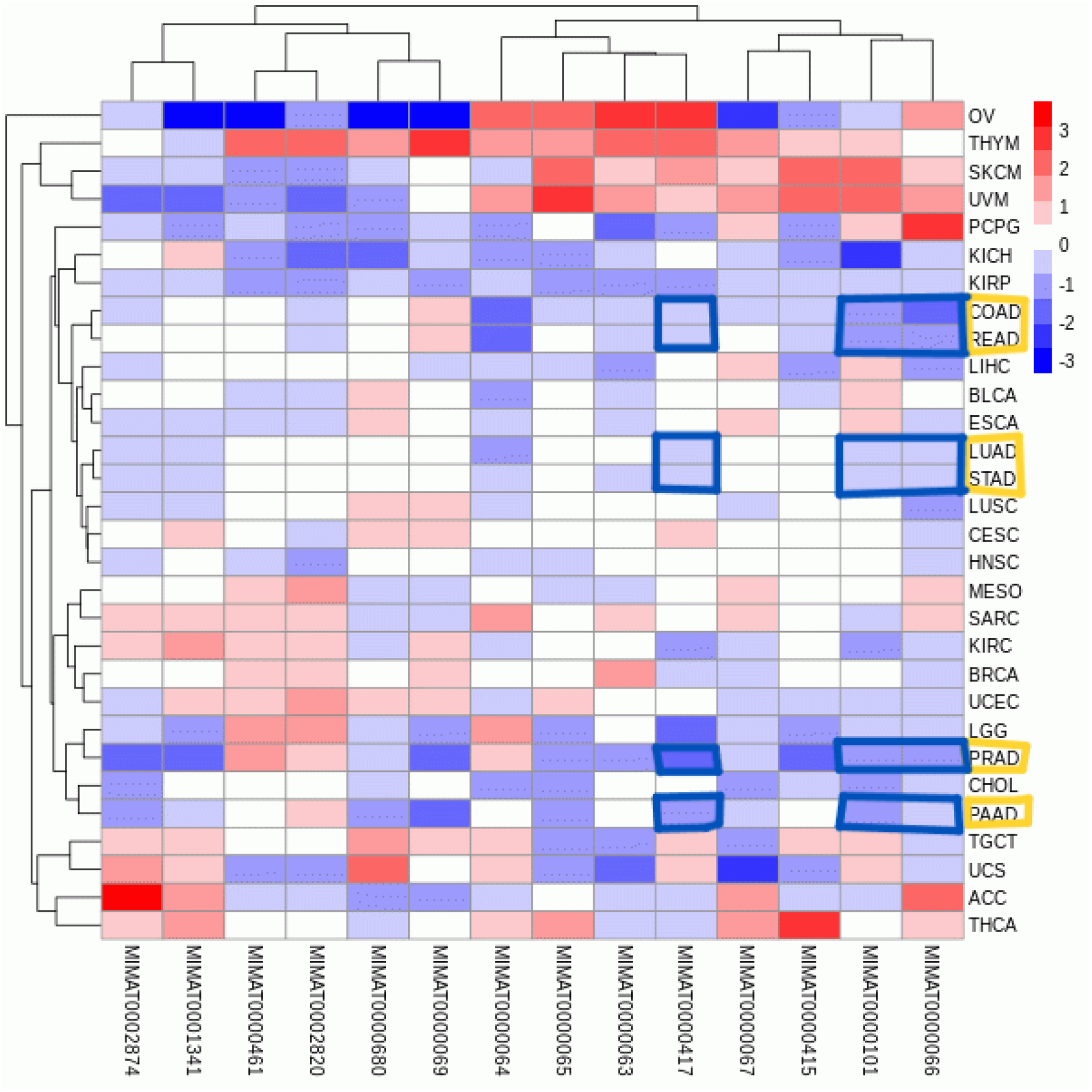
Heatmap of miRNA Expression Across Various Cancer Types. This heatmap visualizes the expression levels of identified miRNAs (labeled with their corresponding MIMAT IDs) across a wide range of cancer types (adenocarcinoma types are boxed in yellow on the right). The color scale represents the relative expression levels, with red indicating up-regulation, blue indicating down-regulation, and white indicating neutral expression levels. Highlighted in blue boxes are specific adenocarcinoma types, including COAD (colon adenocarcinoma), READ (rectum adenocarcinoma), LUAD (lung adenocarcinoma), STAD (stomach adenocarcinoma), PRAD (prostate adenocarcinoma), and PAAD (pancreatic adenocarcinoma). These boxed regions show consistent down-regulation across these adenocarcinoma types of miRNAs hsa-miR-103a-3p, hsa-let-7e-5p, and hsa-miR-15b-5p.The hierarchical clustering along the top and left axes groups the miRNAs and cancer types based on similarity in expression profiles, revealing patterns of miRNA dysregulation that may be characteristic of specific cancer types, including adenocarcinomas. This analysis underscores the potential role of these miRNAs as biomarkers or therapeutic targets in adenocarcinoma.

**Fig. 6:**
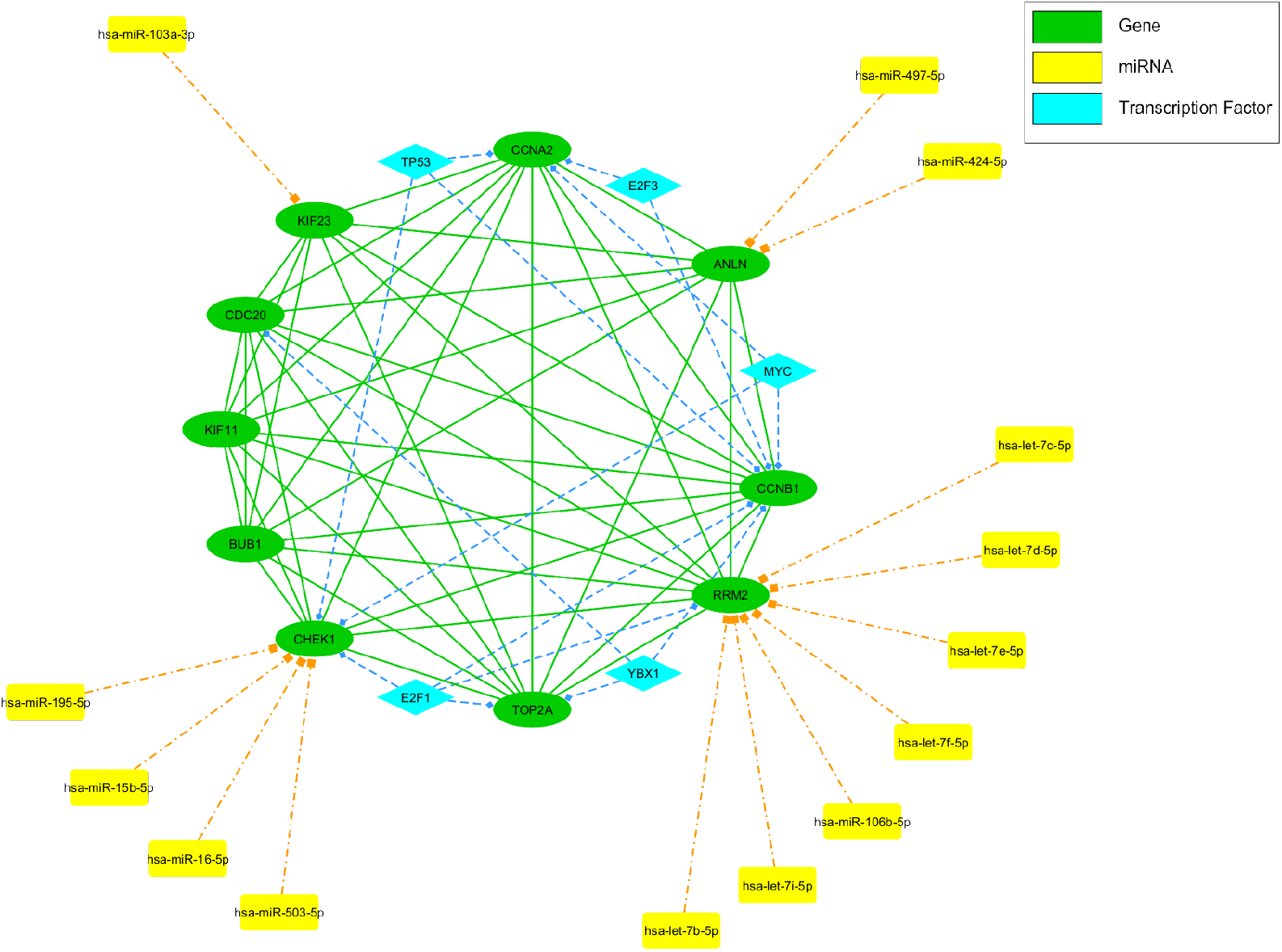
Regulatory Network of Hub Genes, Transcription Factors, and miRNAs in Adenocarcinoma. This figure illustrates the complex regulatory network involving the identified hub genes, transcription factors (TFs), and miRNAs in adenocarcinoma. The network consists of: -**Hub Genes (Green Ovals)**: The core hub genes identified in the study, including CHEK1, CDC20, ANLN, RRM2, CCNB1, CCNA2, KIF23, TOP2A, BUB1, and KIF11. These genes are central to the protein-protein interaction (PPI) network and are significantly overexpressed in various adenocarcinoma types. -**Transcription Factors (Blue Diamonds)**: Key TFs targeting the hub genes, such as YBX1, E2F1, MYC, E2F3, and TP53, are shown to regulate the expression of these critical genes. These TFs play a crucial role in controlling cell cycle progression and other cancer-related processes. - **miRNAs (Yellow Rectangles)**: The miRNAs predicted to target the hub genes include hsa-miR-503-5p, hsa-miR-106b-5p, hsa-miR-16-5p, hsa-miR-103a-3p, hsa-let-7b-5p, hsa-let-7c-5p, hsa-let-7d-5p, hsa-let-7e-5p, hsa-let-7f-5p, hsa-let-7i-5p, hsa-miR-15b-5p, hsa-miR-195-5p, hsa-miR-424-5p, and hsa-miR-497-5p. These miRNAs are shown to interact with multiple hub genes, suggesting their potential role in finetuning the expression of genes critical for adenocarcinoma progression.The network highlights the intricate interactions between these molecular players, where transcription factors regulate the hub genes, and miRNAs further modulate their expression. This multi-layered regulatory framework underscores the complexity of gene regulation in adenocarcinoma and points to potential therapeutic targets for disrupting key pathways involved in tumor growth and progression.

### Functional Enrichment Analysis

Functional enrichment analysis of the 129 common up-regulated genes revealed significant enrichment in several biological processes and pathways relevant to adenocarcinoma. The most prominent categories included cell cycle regulation, mitotic regulation, mitotic nuclear division, cellular senescence, and G1 to S cell cycle control (Figs. 7 and 8). These findings suggest that the identified hub genes play critical roles in the control of cell proliferation and survival, key processes in the pathogenesis of adenocarcinoma.

**Fig. 7:**
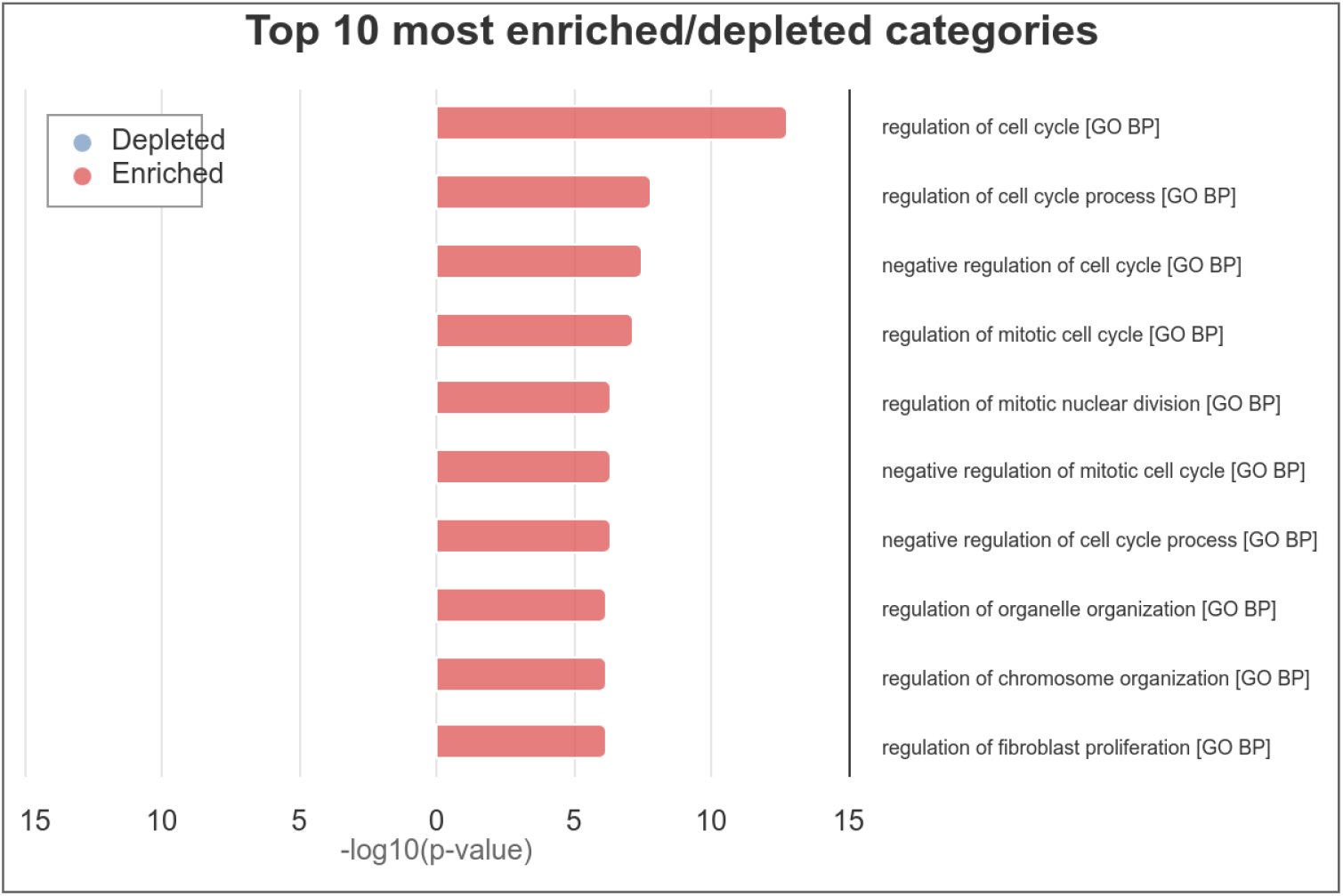
Top 10 Most Enriched Gene Ontology (GO) Biological Process Categories in Up-Regulated Hub Genes in Adenocarcinoma. This bar chart displays the top 10 most significantly enriched Gene Ontology (GO) biological process (BP) categories associated with the up-regulated hub genes and transcription factors identified in adenocarcinoma. The categories are ranked based on the -log10(p-value), indicating the statistical significance of enrichment. **-Enriched Categories (Red Bars)**: The chart highlights that the most enriched biological processes are primarily related to the regulation of the cell cycle, mitotic processes, and cellular organization. Specifically, categories such as “regulation of cell cycle,” “regulation of mitotic cell cycle,” and “regulation of mitotic nuclear division” are strongly represented, reflecting the crucial role of cell cycle control in the pathogenesis of adenocarcinoma. **-Key Insights**: The prominence of these categories underscores the importance of dysregulated cell cycle processes in adenocarcinoma development and progression, suggesting that the identified hub genes play central roles in driving tumor cell proliferation and survival through these pathways. This analysis provides a deeper understanding of the biological functions most affected by the up-regulated genes in adenocarcinoma, offering potential targets for therapeutic intervention.

**Fig. 8:**
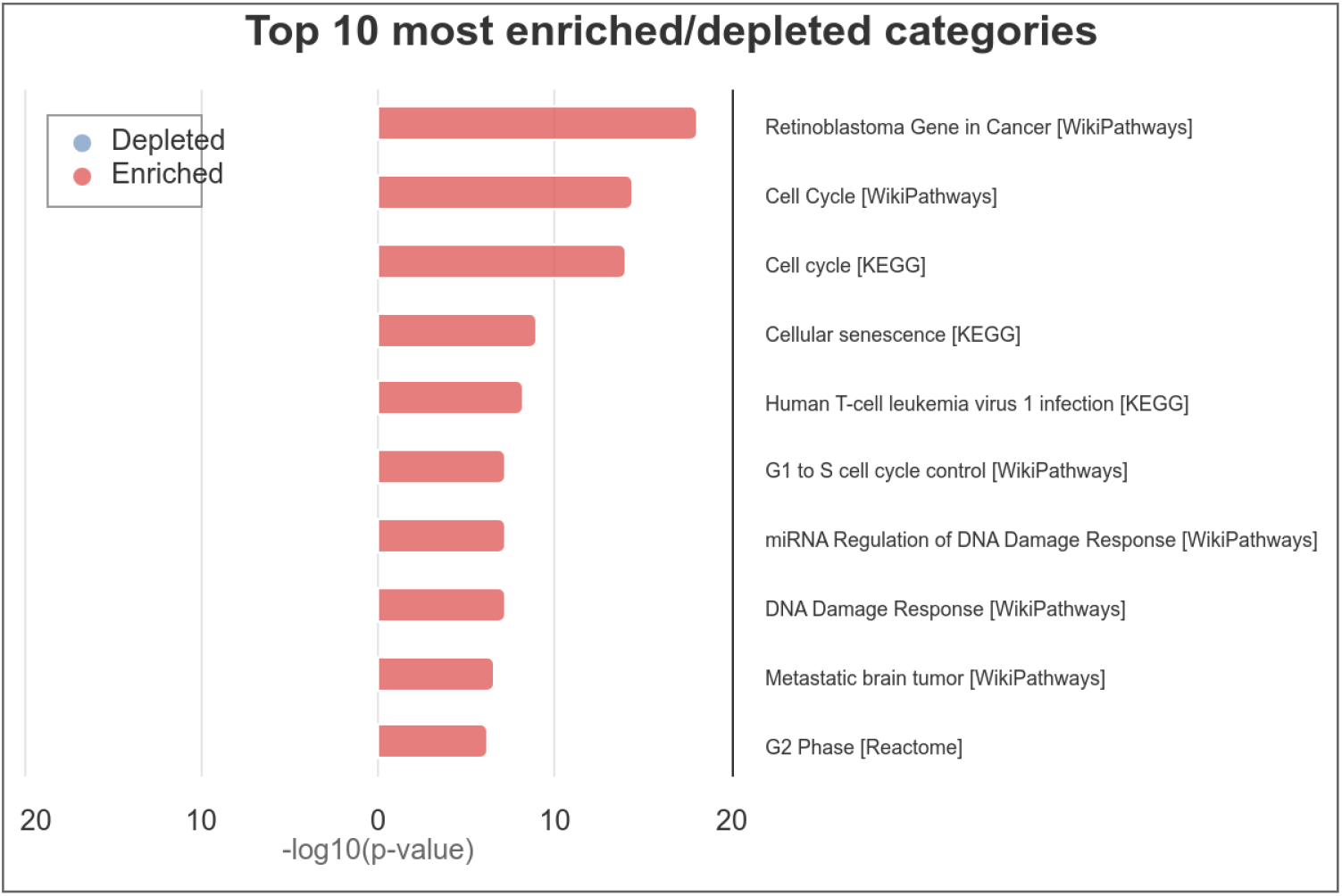
Top 10 Most Enriched Pathway Categories in Up-Regulated Hub Genes in Adenocarcinoma. This bar chart depicts the top 10 most significantly enriched pathway categories associated with the up-regulated hub genes identified in adenocarcinoma. The categories are ranked by the -log10(p-value), reflecting the statistical significance of the enrichment. **- Enriched Categories (Red Bars)**: The analysis reveals that the most enriched pathways are closely related to cell cycle regulation, DNA damage response, and cellular senescence. Specifically, categories such as “Retinoblastoma Gene in Cancer,” “Cell Cycle” (from both WikiPathways and KEGG), “Cellular Senescence,” and “G1 to S cell cycle control” are prominently featured, indicating their critical role in the pathogenesis of adenocarcinoma. -**Key Insights**: These enriched pathways suggest that dysregulation of cell cycle checkpoints, failure of cellular senescence mechanisms, and aberrant DNA damage response are central to the development and progression of adenocarcinoma. The pathways related to the retinoblastoma gene and miRNA regulation of DNA damage response also highlight potential molecular targets for therapeutic intervention in adenocarcinoma.This enrichment analysis underscores the importance of these pathways in driving the malignant behavior of adenocarcinoma cells, offering insights into potential areas for further research and drug development.

These results collectively provide a comprehensive overview of the molecular mechanisms underlying adenocarcinoma, highlighting key regulatory networks and potential targets for therapeutic intervention.

### 4 Discussion and concluding remarks

In this study, we conducted a comprehensive analysis of up-regulated genes across multiple types of adenocarcinoma to identify key hub genes and their associated regulatory networks. Our findings highlight the significant role of cell cycle regulation, mitotic processes, and DNA damage response in the progression of adenocarcinoma, underscoring the potential of these pathways as therapeutic targets.

The identification of ten hub genes (CHEK1, CDC20, ANLN, RRM2, CCNB1, CCNA2, KIF23, TOP2A, BUB1, and KIF11) provides a valuable resource for understanding the molecular drivers of adenocarcinoma. These genes were found to be consistently overexpressed in multiple adenocarcinoma types, suggesting their central role in tumor growth and survival. Functional enrichment analysis revealed their involvement in critical biological processes, including cell cycle regulation, mitotic division, and DNA damage repair, which are known to be dysregulated in cancer.

The regulatory network constructed in this study, incorporating transcription factors such as TP53 and MYC, and miRNAs including hsa-miR-103a-3p and hsa-let-7e-5p, provides further insights into the complex molecular interactions that govern gene expression in adenocarcinoma. The consistent downregulation of certain miRNAs across multiple cancer types highlights their potential as biomarkers for early detection and as targets for therapeutic intervention.

Despite the strengths of our integrative approach, several limitations should be acknowledged. First, the study is based on the analysis of publicly available gene expression datasets, which may have inherent biases due to differences in sample preparation, data processing, and experimental conditions. Additionally, while our findings identify potential therapeutic targets, experimental validation in vitro and in vivo is necessary to confirm the functional roles of the identified hub genes and regulatory elements. It is also important to state that the methodology was applied to down-regulated genes, but no significant ones were found.

Furthermore, the heterogeneity of adenocarcinoma, both between and within different organ sites, presents a challenge for developing universal therapeutic strategies. The identified hub genes may have varying degrees of relevance in different adenocarcinoma subtypes, which underscores the need for personalized approaches in cancer treatment.

Future research should focus on validating the biological significance of these hub genes and their associated regulatory networks in adenocarcinoma. In particular, the development of targeted therapies that can modulate the activity of these genes or their regulatory elements holds promise for improving patient outcomes. Additionally, integrating gene expression data with other omics approaches, such as proteomics and metabolomics, could provide a more comprehensive understanding of adenocarcinoma biology and uncover novel therapeutic opportunities.

In conclusion, this study enhances our understanding of the molecular mechanisms underlying adenocarcinoma and identifies key regulatory networks that could serve as potential therapeutic targets. These findings lay the groundwork for future investigations aimed at developing more effective and personalized treatment strategies for adenocarcinoma patients.

## Supporting information

Supplemental boxplots

## Author contributions

**Ricardo Romero:** Conceptualization (Lead), Data curation (Equal), Formal analysis (Equal), Investigation (Lead), Methodology (Lead), Writing – original draft (Lead), Writing – review \& editing (Lead).

**Dulce Bastida:** Data curation (Equal), Formal analysis (Equal), Investigation (Supporting), Methodology (Supporting), Writing – review \& editing (Supporting).

## Data availability statement

The data that support the findings of this study are openly available in public databases cited in the text.

## References

[1] Bray F, Ferlay J, Soerjomataram I, Siegel RL, Torre LA, Jemal A. Global cancer statistics 2018: GLOBOCAN estimates of incidence and mortality worldwide for 36 cancers in 185 countries. CA Cancer J. Clin. 2018;68(6):394–394.

[2] Travis WD, Brambilla E, Noguchi M, et al. International association for the study of lung cancer/american thoracic society/european respiratory society international multidisciplinary classification of lung adenocarcinoma. J. Thorac. Oncol. 2011;6(2):244–244.

[3] Brenner H, Kloor M, Pox CP. Colorectal cancer. Lancet. 2014;383(9927):1490–1490.

[4] Siegel RL, Miller KD, Jemal A. Cancer statistics, 2020. CA Cancer J. Clin. 2020;70(1):7–7.

[5] Hanahan D, Weinberg RA. Hallmarks of cancer: the next generation. Cell. 2011;144(5):646–646.

[6] Cancer Genome Atlas Research Network. Comprehensive molecular profiling of lung adenocarcinoma. Nature. 2014;511(7511):543–543.

[7] Vogelstein B, Papadopoulos N, Velculescu VE, Zhou S, Diaz LA, Kinzler KW. Cancer genome landscapes. Science. 2013;339(6127):1546–1546.

[8] Sanchez-Vega F, Mina M, Armenia J, et al. Oncogenic signaling pathways in The Cancer Genome Atlas. Cell. 2018;173(2):321–321.e10.

[9] Cieślik M, Chinnaiyan AM. Cancer transcriptome profiling at the juncture of clinical translation. Nat. Rev. Genet. 2018;19(2):93–93.

[10] Mok TS, Wu YL, Thongprasert S, et al. Gefitinib or carboplatin-paclitaxel in pulmonary adenocarcinoma. N. Engl. J. Med. 2009;361(10):947–947.

[11] Kopetz S, Grothey A, Yaeger R, et al. Encorafenib, binimetinib, and cetuximab in BRAF V600E-mutated colorectal cancer. N. Engl. J. Med. 2019;381(17):1632–1632.

[12] Dagogo-Jack I, Shaw AT. Tumour heterogeneity and resistance to cancer therapies. Nat. Rev. Clin. Oncol. 2018;15(2):81–81.

[13] Vasan N, Baselga J, Hyman DM. A view on drug resistance in cancer. Nature. 2019;575(7782):299–309.

[14] Pereira SP, Oldfield L, Ney A, et al. Early detection of pancreatic cancer. Lancet Gastroenterol. Hepatol. 2020;5(7):698–698.

[15] Quail DF, Joyce JA. Microenvironmental regulation of tumor progression and metastasis. Nat. Med. 2013;19(11):1423–1423.

[16] Slack FJ, Chinnaiyan AM. The role of non-coding RNAs in oncology. Cell. 2019;179(5):1033–1033.

[17] Hasin Y, Seldin M, Lusis A. Multi-omics approaches to disease. Genome Biol. 2017;18(1):83.

[18] Selamat SA, Chung BS, Girard L, et al. Genome-scale analysis of DNA methylation in lung adenocarcinoma and integration with mRNA expression. Genome Res. 2012;22(7):1197–1197.

[19] Jiang X, Tan J, Li J, et al. DACT3 is an epigenetic regulator of Wnt/beta-catenin signaling in colorectal cancer and is a therapeutic target of histone modifications. Cancer Cell. 2008;13(6):529– 541.

[20] Badea L, Herlea V, Dima SO, Dumitrascu T, Popescu I. Combined gene expression analysis of whole-tissue and microdissected pancreatic ductal adenocarcinoma identifies genes specifically overexpressed in tumor epithelia. Hepatogastroenterology. 2008;55(88):2016–2016.

[21] Idichi T, Seki N, Kurahara H, et al. Regulation of actin-binding protein ANLN by antitumor miR-217 inhibits cancer cell aggressiveness in pancreatic ductal adenocarcinoma. Oncotarget. 2017;8(32):53180–53180.

[22] Tang Z, Kang B, Li C, Chen T, Zhang Z. GEPIA2: an enhanced web server for large-scale expression profiling and interactive analysis. Nucleic Acids Res. 2019;47(W1):W556–W560.

[23] Szklarczyk D, Kirsch R, Koutrouli M, et al. The STRING database in 2023: protein-protein association networks and functional enrichment analyses for any sequenced genome of interest. Nucleic Acids Res. 2023;51(D1):D638–D646.

[24] Shannon P, Markiel A, Ozier O, et al. Cytoscape: a software environment for integrated models of biomolecular interaction networks. Genome Res. 2003;13(11):2498–2498.

[25] Chin CH, Chen SH, Wu HH, Ho CW, Ko MT, Lin CY. cytoHubba: identifying hub objects and sub-networks from complex interactome. BMC Syst. Biol. 2014;8 Suppl 4(S4):S11.

[26] Sticht C, De La Torre C, Parveen A, Gretz N. miRWalk: An online resource for prediction of microRNA binding sites. PLoS One. 2018;13(10):e0206239.

[27] McGeary SE, Lin KS, Shi CY, et al. The biochemical basis of microRNA targeting efficacy. Science. 2019;366(6472):eaav1741.

[28] Chen Y, Wang X. miRDB: an online database for prediction of functional microRNA targets. Nucleic Acids Res. 2020;48(D1):D127–D131.

[29] Huang HY, Lin YCD, Cui S, et al. miRTarBase update 2022: an informative resource for experimentally validated miRNA-target interactions. Nucleic Acids Res. 2022;50(D1):D222–D230.

[30] Wong NW, Chen Y, Chen S, Wang X. OncomiR: an online resource for exploring pan-cancer microRNA dysregulation. Bioinformatics. 2018;34(4):713–713.

[31] Han H, Cho JW, Lee S, et al. TRRUST v2: an expanded reference database of human and mouse transcriptional regulatory interactions. Nucleic Acids Res. 2018;46(D1):D380–D386.

[32] Gerstner N, Kehl T, Lenhof K, et al. GeneTrail 3: advanced high-throughput enrichment analysis. Nucleic Acids Res. 2020;48(W1):W515–W520.

[33] Ashburner M, Ball CA, Blake JA, et al. Gene ontology: tool for the unification of biology. The Gene Ontology Consortium. Nat. Genet. 2000;25(1):25–25.

[34] Consortium GO, Aleksander SA, Balhoff J, et al. The Gene Ontology knowledgebase in 2023. Genetics. 2023;224(1).

