## Supplemental boxplots for "Identification and Functional Characterization of Up-Regulated Hub Genes in Adenocarcinoma Across Multiple Organ Sites"

Expression level of Tumor (in salmon) vs Normal (in grey) samples of hub genes across seven adenocarcinoma types (COAD, LUAD, PAAD, PRAD, READ, STAD, and OV), obtained from GEPIA2. The red asterisk denotes significance (adjusted P value < 0.05, |logFC| > 1)

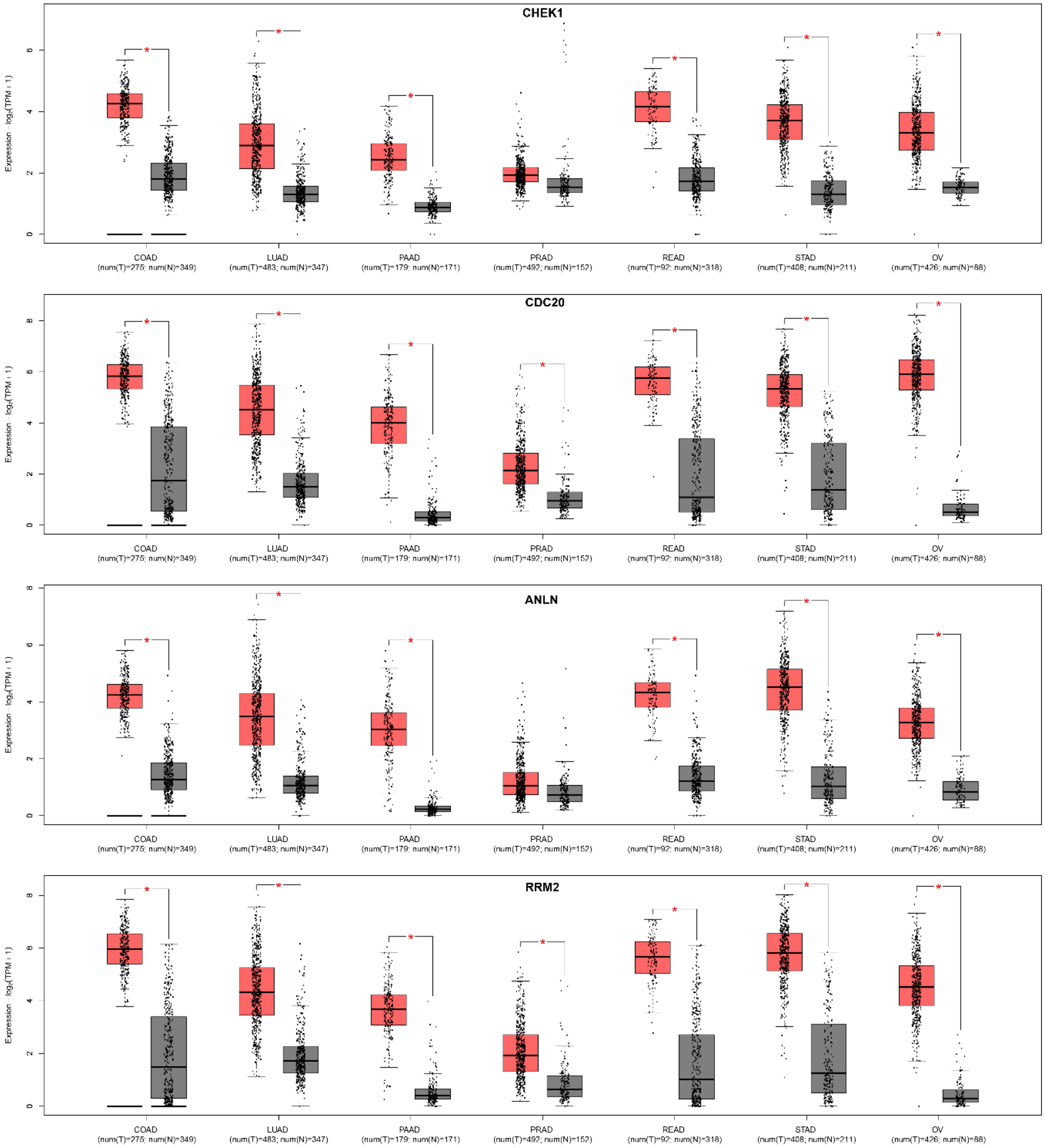

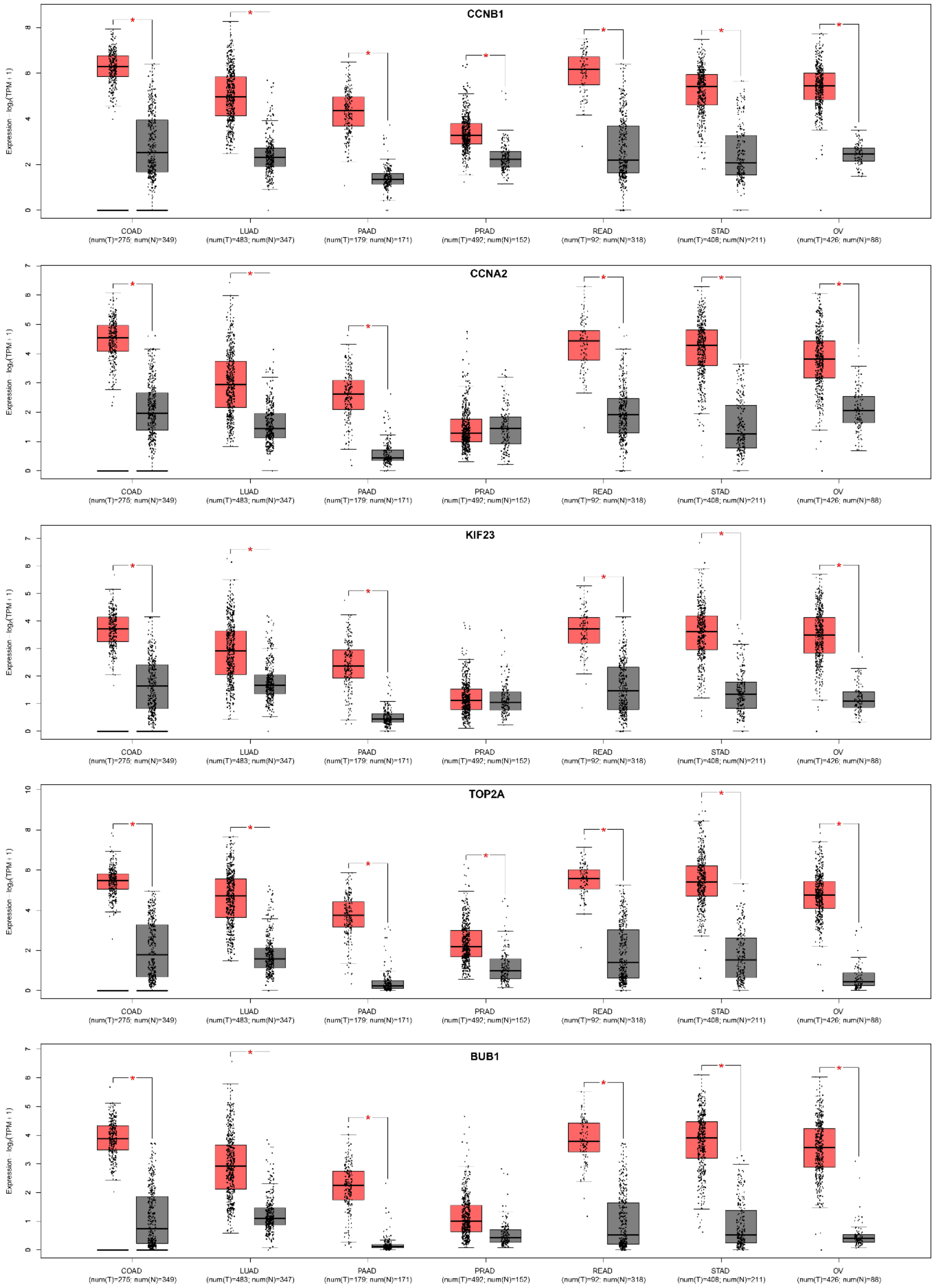
CCNB1
CCNA2
KIF23
TOP2A
BUB1
Expression - log2(TPM+1)
COAD (num(T)=275; num(N)=349)
LUAD (num(T)=483; num(N)=347)
PAAD (num(T)=179; num(N)=171)
PRAD (num(T)=492; num(N)=152)
READ (num(T)=92; num(N)=318)
STAD (num(T)=408; num(N)=211)
OV (num(T)=426; num(N)=88)

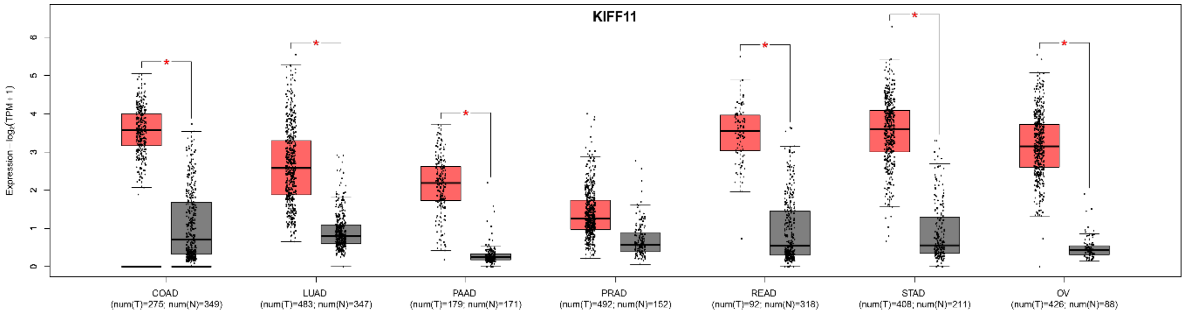
KIFF11
Expression – log2(TPM + 1)
0
1
2
3
4
5
6
*
COAD
(num(T)=275; num(N)=349)
LUAD
(num(T)=483; num(N)=347)
PAAD
(num(T)=179; num(N)=171)
PRAD
(num(T)=492; num(N)=152)
READ
(num(T)=92; num(N)=318)
STAD
(num(T)=408; num(N)=211)
OV
(num(T)=426; num(N)=88)
